# Identification and quantification of neurological responses in patients with dentine hypersensitivity

**DOI:** 10.64898/2026.06.29.735173

**Authors:** Natalie Wong, Hannah I Barnes, Charles R Parkinson, Mark W Barber, Mahnaz Arvaneh, Fiona M Boissonade

## Abstract

Evaluation of the effectiveness of therapeutic interventions for dentine hypersensitivity is limited by a lack of standardisation and objectivity in measuring the associated pain. To address this, we investigated whether electroencephalography (EEG) can provide an objective, quantitative measure of the condition. Participants with and without dentine hypersensitivity underwent evaporative (air puff) and thermal (cooling probe) tooth stimulation during continuous recording of EEG activity. Sensitivity scores (Schiff Sensitivity score for air puff stimuli, and Visual Analogue Scale score (VAS) for thermal stimuli) were recorded, and participants’ responses to the Dentine Hypersensitivity Experience Questionnaire (DHEQ) collected.

There were strong positive correlations between the Schiff and VAS scores, and also between both sensitivity scores and the impact of dentine hypersensitivity on quality of life (DHEQ). Additionally, EEG data analysis revealed significant differences in event-related potentials (ERP) following evaporative stimulation between participants with different Schiff scores, and in cortical activity between traces where participants indicated discomfort and those where participants did not indicate discomfort during thermal stimulation trials.

Topographical maps of EEG band power during thermal stimulation showed progressive cortical recruitment and focal activation emerging in the 3 seconds prior to indication of discomfort. Comparison of EEG band power between response and no response trials to thermal stimulation showed significantly higher delta frequency band power in response trials than in no-response trials.

Peak-to-peak amplitude of cortical response during thermal stimulation correlated with DHEQ and VAS scores, and the probe temperature at which participants indicated discomfort.

These findings suggest that components of EEG responses align with other measures of dentine sensitivity (DHEQ, Schiff and VAS scores) and can serve as objective neurophysiological markers for evaluating the severity of dentine hypersensitivity.

## Introduction

Dentine hypersensitivity is a common dental condition that has been defined as a ‘short, sharp pain arising from exposed dentine in response to stimuli typically thermal, evaporative, tactile, osmotic or chemical and which cannot be ascribed to any other form of dental defect or pathology’ [1]. Many interventions for dentine hypersensitivity have been investigated over the last 60 years, but the lack of standardisation and objectivity in measuring pain associated with dentine hypersensitivity are major limitations for assessing the efficacy of various therapeutic interventions. Consequently, there is a need to develop quantitative, objective markers that can complement traditional subjective diagnostic tools.

Electroencephalography (EEG) is a non-invasive technique that measures the electrical activity of the brain using electrodes placed on the scalp. It offers a highly sensitive method with good temporal resolution to investigate the brain’s real-time response to noxious stimuli [2,3]. Within EEG analysis, event-related potentials (ERPs) are time-locked electrical responses that are associated with specific sensory, cognitive, and motor events [4]. By analysing ERPs, the brain’s response to air and thermal stimuli applied to the tooth can be isolated. Therefore, this study aims to investigate whether EEG responses to tooth stimulation can be used to evaluate the severity of dentine hypersensitivity objectively. Furthermore, this study aims to explore whether participants’ responses to thermal stimulation, measured using the visual analogue scale (VAS) score, can be used to assess dentine hypersensitivity. Such understanding would provide a significant step forward for the quantitative assessment of the efficacy of therapeutic interventions directed towards alleviating dentine hypersensitivity.

## Methods

The study explored and compared EEG responses to tooth stimulation in participants with and without dentine hypersensitivity. Two methods of sensitivity stimulation were used to explore if different stimuli resulted in different EEG responses: evaporative stimulation using a dental triple syringe and thermal stimulation using an intra-oral thermal probe. A schematic overview of the study design is presented in Figure 1.

**Figure 1.**
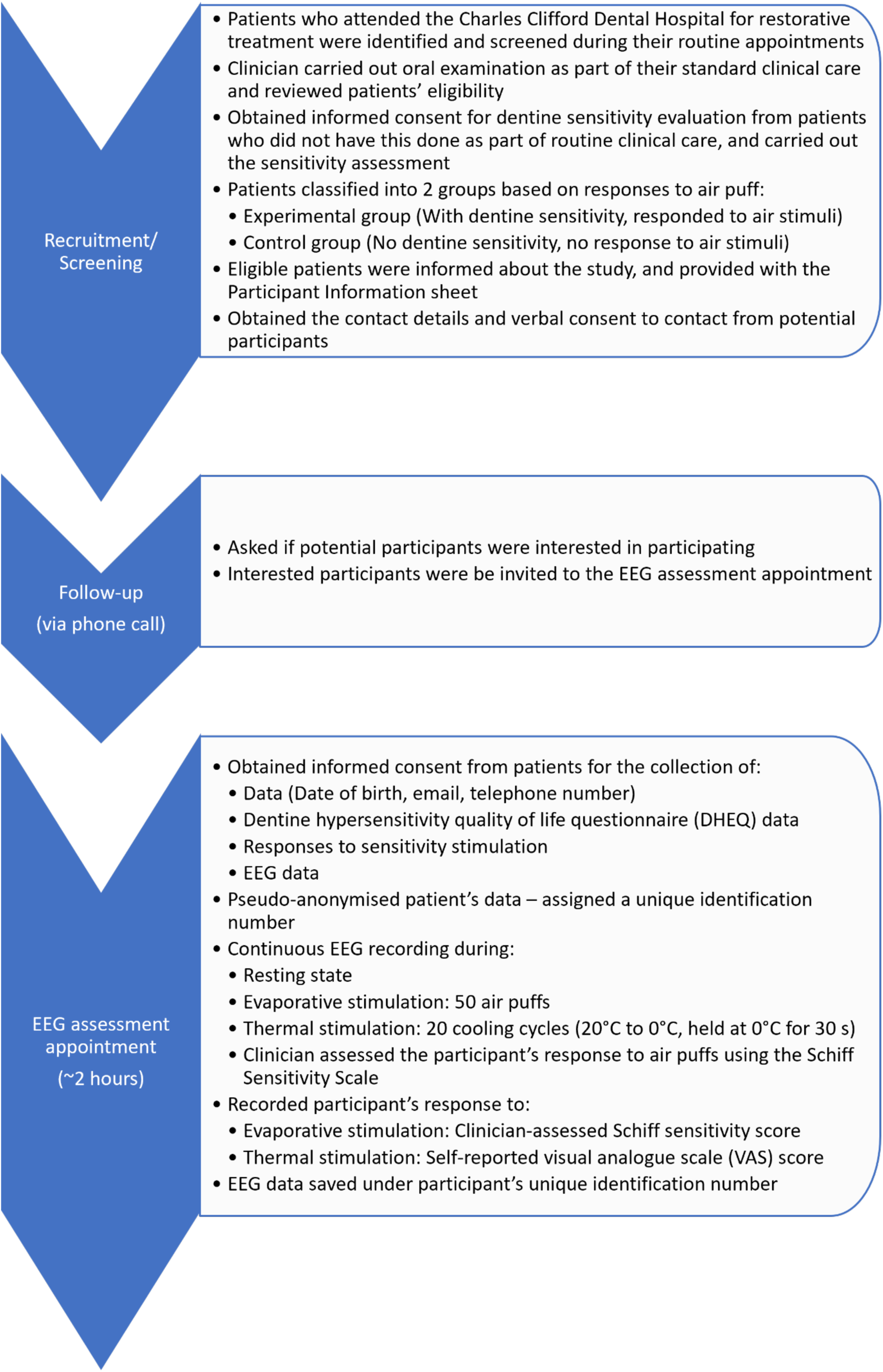
Schematic overview of the study design.

Following screening, participants were classified into the experimental (with dentine hypersensitivity; n=30) and control (no dentine hypersensitivity; n=30) groups based on their responses to air puff stimuli. Data were pseudo-anonymised by assigning each participant a unique identification number. Participants completed the Dentine Hypersensitivity Experience Questionnaire (DHEQ) to assess the impact of dentine hypersensitivity on quality of life. Continuous EEG signals were recorded during resting state and throughout the two methods of sensitivity stimulation (evaporative and thermal). Finally, their responses to evaporative and thermal stimulation were assessed using the clinician-assessed Schiff Sensitivity Scale and the self-reported visual analogue scale (VAS) score respectively.

### Participants

60 adults (26 males, 34 females) between the ages of 18-64 were recruited for the study. All participants were recruited with their written informed consent, in accordance with ethical approvals from the NHS Health Research Authority (HRA) and Sheffield Teaching Hospitals (STH) (24/YH/0159 STH22405). To pseudo-anonymise the data, each participant was assigned a unique identification number, and participants were only identified by their identification number on all data collection documents.

Patients attending routine restorative appointments at the Charles Clifford Dental Hospital were screened as potentially eligible participants for the study by members of the direct clinical care team. The inclusion criteria of the study required participants to be at least 18 years of age, in good general health, and possess a minimum of 10 teeth (excluding those with crowns or bridges) distributed from the right to the left first premolars in both the upper and lower arches. Dental exclusion criteria included the use of maxillary or mandibular orthodontic appliances, obvious signs of untreated caries, evidence of periodontitis, or periodontal pocket depths that were 4 mm or greater in the anterior sextants. Participants were also excluded if they had a history of seizures, had damaged scalp skin (e.g., cuts, psoriasis, eczema), or were taking medications known to affect brain responses. Participants were also excluded if they had difficulty understanding English or were unable to comply with the study protocol.

For patients with self-reported dentine hypersensitivity, the direct clinical care team evaluated their tooth sensitivity via responses to air puffs as part of their standard clinical care. For patients presenting with no self-reported dentine hypersensitivity, written informed consent was obtained to perform dentine hypersensitivity evaluation. Their teeth were tested with air puffs to examine if they have dentine hypersensitivity. Based on their response to the air puff stimuli, 30 participants were classified into the experimental group (with dentine hypersensitivity) and 30 into the control group (no dentine hypersensitivity). One tooth per participant was selected for testing. For the experimental group, the test tooth was selected based on responses to air puff. For the control group, the test tooth was selected to match the location of a corresponding tooth in the experimental group. To assess the impact of dentine hypersensitivity, all participants were asked to complete the Dentine Hypersensitivity Experience Questionnaire (DHEQ) [5].

### Hardware and interfaces

#### Electroencephalography system

Electroencephalography (EEG) signals were continuously recorded during tooth stimulation using an Enobio 8 system (Neuroelectrics, Spain) via a USB connection to a recording laptop running the Neuroelectrics Instrument Controller (NIC2) software, the EEG system’s software interface. Data were acquired at a sampling rate of 500 Hz from eight channels, categorised by their brain regions: Frontal (F): F7, Fz, F8; Central (C): C3, Cz, C4; and Central Parietal (CP): CP5, CP6, which were positioned according to the 10-20 standard system [6], shown in Figure 2. The electrodes were filled with a salt-based conductive gel to form a link between the electrodes and the participant’s scalp, and were secured within a cap. The Common Mode Sense (CMS) and Driven Right Leg (DRL) electrodes were placed on the right mastoid to serve as the reference. EEG signal quality was monitored via NIC2, and trials only started when all channels were displayed as ‘green’, signifying low impedance and high signal quality.

**Figure 2.**
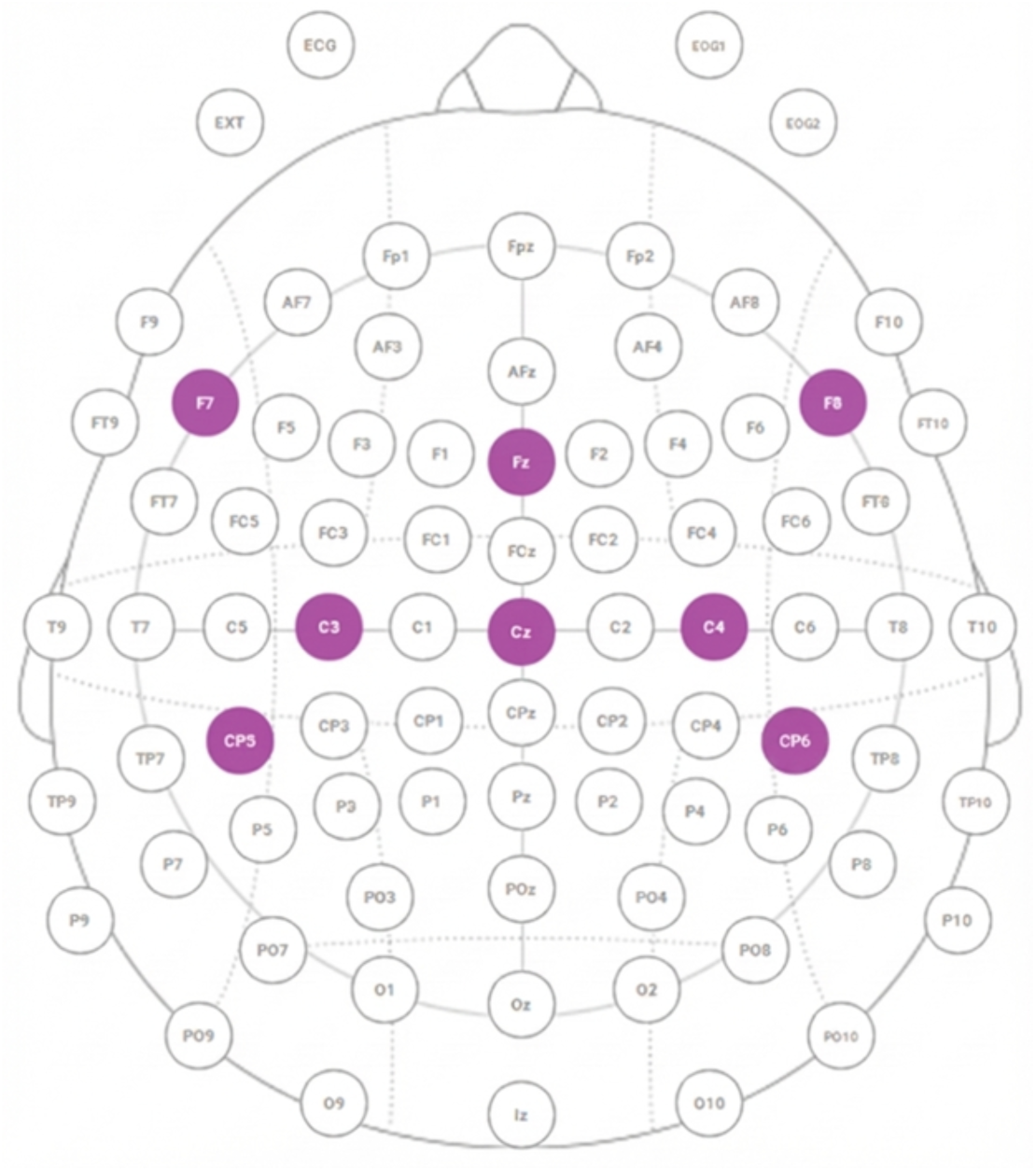
Schematic map of EEG electrode locations. The eight channels used to collect EEG signals in the study are highlighted in purple, and correspond to the Frontal (F7, Fz, F8), Central (C3, Cz, C4), and Central Parietal (CP5, CP6) regions.

#### Tooth stimulation systems and software

Evaporative stimulation was applied using a dental triple syringe (A-dec, United Kingdom) to deliver 1-second puffs of air to the test tooth. Stimulus timing markers were acquired using a push button that was attached to the triple syringe button (when depressed initiates air flow), and wired to a microcontroller (Arduino Nano ESP32). The microcontroller transmitted changes in the state of the push button via WiFi to a MATLAB script on the recording laptop, which then sent the markers to NIC2. This ensured synchronisation between the acquired timing markers and the continuous EEG data. The measured latency between button press and the registration of the marker on NIC2 was 12 ms with a jitter of ±3 ms.

Thermal stimulation was applied to the same test tooth using a computer-based Thermal Sensory Analyzer (TSA) II system (Medoc, Israel), which included an intra-oral thermal probe and the associated Medoc Main Station software that controlled the cooling cycle. The spacebar on the laptop keyboard was used by the experimenter to control the start of cooling cycle. Participants were given an external keyboard and were instructed to press ‘s’ if they felt any discomfort, which aborted the cooling cycle. If the participant did not respond, the experimenter pressed ‘e’ on the laptop keyboard to mark the end of the full cooling cycle. These keyboard presses were continuously monitored by a MATLAB script. Upon detection of a keyboard press (Spacebar, ‘s’, or ‘e’), the script immediately sent a marker to NIC2. This ensured synchronisation between the onset and termination of the thermal stimulus, the participant’s response, and the continuous EEG data.

### Experimental procedure

The study took place at the Charles Clifford Dental Hospital (CCDH) in Sheffield, and was conducted in a surgery room within CCDH. Upon arrival and informed consent, participants were fitted with the EEG cap. They were comfortably seated in a reclined dental chair, with their head resting on the headrest. To ensure high EEG signal quality, participants were asked to remain as still as possible during all periods of tooth stimulation.

The total duration of the study session was approximately 2 hours. During the setup of the EEG cap, participants completed the DHEQ. Data collection involved two sequential methods of sensitivity assessment: evaporative stimulation and thermal stimulation. Participants were free to take a break in between each assessment block.

#### Evaporative stimulation

50 air puffs were applied (using a triple syringe) to the buccal surface of the test tooth, with randomised interstimulus intervals between 3 and 8 seconds. The participant’s response to the air puffs was recorded by the clinical dental professional using the Schiff Sensitivity Scale (0 = participant does not respond to air stimulus; 1 = participant responds to air stimulus but does not request discontinuation; 2 = participant responds to air stimulus and requests discontinuation or moves from stimulus; 3 = participant responds to air stimulus, considers stimulus to be painful and requests discontinuation of the stimulus) [7].

#### Thermal stimulation

The intra-oral thermal probe was placed on the buccal surface of the test tooth. First, First, to determine which elements of the EEG signal were arising from pressing ‘s’ to respond, participants completed five button press control trials. In these trials, participants were asked to intentionally press ‘s’ approximately 10 to 20 seconds after the probe was placed on their tooth, while the probe was held at baseline temperature of 20°C.

The cooling cycle consisted of a baseline temperature of 20°C (held for 3 seconds), followed by a reduction in temperature from 20°C to 0°C at a rate of -1°C/s. The probe was then held at 0°C for 30 seconds. This cycle was repeated 20 times, with a minimum break of 30 seconds in between each cycle. Participants were asked to press ‘s’ on the keyboard that aborted the cooling cycle immediately if they felt any discomfort associated with the thermal stimuli. At this point the thermal probe was removed from the tooth. If no response was given, the thermal probe was removed at the end of cooling cycle. After all cooling cycles were completed, participants were asked to complete a visual analogue scale (VAS) score to indicate the intensity of the sensation they felt. This consisted of a 10 cm straight line with the two endpoints representing 0 (‘I did not feel anything at all’) and 10 (‘It is extremely painful’). The point at which the perpendicular line drawn by the participant intersected the 10 cm line was measured to the nearest millimetre, giving a reading between zero and ten.

### Data analyses

#### Pre-processing of EEG data

Due to poor quality of data collected, EEG data from one participant in the control group were excluded from further analysis. A total of 59 participants (29 in the control group and 30 in the experimental group) were included in the EEG data analysis. All EEG data processing was performed using MATLAB R2023b. EEG data were first filtered by applying a zero-phase, 16^th^-order Butterworth bandpass filter with cutoff frequencies set between 0.5 Hz and 40 Hz. Artefact rejection was then performed using the moving-window technique [4], by excluding data segments where the difference between the minimum and maximum amplitude within a 400-sample window exceeded the threshold of 50 µV in any individual channel. Segments failing this criterion were marked for exclusion.

#### Analysis of evaporative stimulation data

##### Analysis of evaporative stimulus timing markers

Due to a technical error in the initial Arduino code, only the air puff offset markers were logged for the first 48 participants. For the remaining 11 participants, both the onset and offset markers were successfully recorded. To calculate the missing air puff onset times, the air puff duration (Duration_est_) was conservatively estimated using the measured durations collected from the 11 participants (Duration_11 Ps_): Duration_est_= Median(Duration_11 Ps_) − 2 × SD(Duration_11 Ps_), which yielded an estimated duration of 1070 ms. The estimated air puff onset was then calculated as Time_offset_ − 1070 ms, which was applied when analysing data from all 59 participants to maintain consistency across all participants.

##### Epoching and baseline correction

Following initial EEG data filtering and artefact rejection, data were epoched relative to the recorded air puff offset markers. Based on the estimated air puff duration, each trial was epoched from 1070 ms before to 1000 ms after the offset. The pre-stimulus baseline was defined as the 100 ms window immediately preceding the estimated air puff onset, which corresponded to the interval from -1170 ms to -1070 ms relative to the offset marker. Baseline correction was applied by subtracting the mean amplitude within this 100 ms baseline window from each channel and trial.

##### Data aggregation and statistical analysis

The analysis focused on four channels: C3, C4, Cz, and Fz, which were selected based on the predicted cortical involvement of the somatosensory and motor systems. C3 and C4 capture activity from the primary somatosensory cortex [8], which processes sensory information from the body. These channels were dynamically assigned as ipsilateral or contralateral based on the side of the test tooth. Specifically, for a test tooth on the right side, C4 was designated as ipsilateral and C3 as contralateral; for a test tooth on the left side, C3 was designated as ipsilateral and C4 as contralateral. Cz is primarily associated with the primary motor cortex [9], which is responsible for controlling and executing movements. Fz is located over the central frontal region, capturing activity related to motor planning, as well as attention and cognitive control [10].

All valid trials were aggregated based on the participant’s Schiff score (0, 1, or 2). The extracted epochs were analysed using a mass univariate, sliding window approach on the three Schiff score groups. This involved calculating the mean amplitude within sequential 100 ms time windows (overlapping by 50 ms) for each group. These mean amplitudes were then used to perform pairwise comparisons (Schiff score = 0 vs 1, Schiff score = 0 vs 2, and Schiff score = 1 vs 2) using Welch’s t-test. Bonferroni correction was applied to control the family-wise error rate across the three pairwise tests per time window, resulting in a corrected significance threshold of α = 0.0167.

For visualisation, the mean responses to evaporative stimulation (the time window of 1070 ms before to 1000 ms after the air puff offset) and their corresponding standard error of the mean (SEM) traces were calculated for all three Schiff score groups across the four channels. A zero-phase, 4th-order Butterworth low-pass filter with a cutoff frequency of 8 Hz was then applied to smooth both traces prior to plotting. Time windows with statistically significant difference between groups, as determined by the Bonferroni-corrected Welch’s t-tests, were indicated by shading on the figures.

#### Analysis of thermal stimulation data

##### Epoching, baseline correction and button press response correction

Following initial EEG data filtering and artefact rejection, EEG data corresponding to the cooling cycles were segmented based on trial outcome: response and no response trials. The analysis window for all trials spanned a total of 4 seconds (-3000 ms to +1000 ms) relative to the trial’s termination marker. Specifically, for response trials, data were epoched relative to the time of participant’s response; for no response trials, data were epoched relative to the experimenter-marked end of cooling cycle.

The pre-stimulus baseline window was defined as the 2-second window that began 1 second after the initial start marker (when the probe was held at 20°C). Baseline correction was applied by calculating the mean amplitude within this 2-second baseline window and subtracting this mean from the corresponding 4-second analysis window.

To eliminate responses related to the action of button pressing, cortical activity associated with this activity (recorded during the button press control trials, described in section 3.2) was subtracted from the analysis windows of response trials. For each participant, a button press response template was constructed by averaging the same 4-second epochs (-3000 ms to +1000 ms) relative to the trial’s termination marker of their button press control trials. This individual-specific button press response template was then subtracted from the analysis windows of the corresponding participant’s response trials. This ensured that the final response group epochs used for subsequent analysis activity related to nociceptive processing, rather than the motor execution of button press.

##### Data aggregation and statistical analysis

The analysis focused on the same four channels established in the evaporative stimulation data analysis: C3, C4, Cz, and Fz. As described previously, the C3 and C4 channels were dynamically assigned as ipsilateral or contralateral based on the side of the test tooth for subsequent analysis.

All valid trials were aggregated into two outcome groups: response and no response. The extracted epochs were analysed using a mass univariate, sliding window approach. This involved calculating the mean amplitude within sequential 100 ms time windows (overlapping by 50 ms) for both groups. Welch’s t-test was performed at each time window to compare the mean amplitudes between the response and no response groups. A significance threshold of p < 0.05 was applied.

For visualisation, the mean responses and their corresponding SEM traces, calculated across the analysis window, were plotted for both response and no response groups across the four channels. A zero-phase, 4th-order Butterworth low-pass filter with a cutoff frequency of 8 Hz was then applied to smooth both traces prior to plotting. Time windows with statistically significant difference between groups were indicated by grey shading on the figures.

##### Topographical analysis

Topographical maps were generated to visualise the spatial distribution of band power (detailed below) across the scalp. The data used for plotting first underwent baseline correction and button press response removal. The maps were plotted using the overall mean of participant-averaged band power for sequential time segments, covering the 5-second period before the trial termination marker (participant’s response or end of cooling cycle). Data from all eight available channels were used to generate the maps for the ten consecutive 0.5 s segments. Separate figures were generated by grouping trials based on the side of the test tooth (right side or left side) to visualise the differences between response and no response groups.

##### Band power quantification

After baseline correction and button press response removal (for response trials only), time-frequency analysis was performed to quantify band power across different frequency bands: delta (1-4 Hz), theta (4-8 Hz), alpha (8-12 Hz), beta (12-30 Hz), and gamma (30-100 Hz). Spectral power was computed for the 2-second time window (-2000 ms to 0 ms) prior to the trial’s termination marker, a window selected to capture neural activity preceding the decision to indicate discomfort. The mean band power for each channel and frequency band was calculated per participant for both response and no response groups. To compare band power differences between these two groups, either an unpaired t-test or the non-parametric Mann-Whitney test was used, which was determined after performing the Shapiro-Wilk test to assess the normality of data.

#### Analysis of EEG and behavioural markers

The correlation between DHEQ scores and sensitivity scores (Schiff score and VAS score) was assessed using the Spearman’s rank correlation.

##### Peak-to-peak amplitude quantification and correlation

For the purpose of correlational analysis with various behavioural markers, all valid trials were first processed: baseline correction was applied, and button press response removal was completed for response trials. Key peak components were then identified by averaging all valid, processed trials (combining response and no response trials) for each participant.

The negative peak was identified by locating the minimum amplitude within the time window spanning -1000 ms to -500 ms relative to trial termination, whereas the positive peak was identified by locating the maximum amplitude within the time window spanning -300 ms to +200 ms relative to trial termination. The peak-to-peak amplitude (calculated independently for all four channels) was then calculated as the difference between positive peak and negative peak amplitude.

The temperature at which the participant pressed ‘s’ was recorded for cooling cycles with response. To account for cooling cycles with no response, the temperature of response for those trials was set as -5°C (the technical limit of the thermal probe). The adjusted temperature of response for each participant was then calculated by averaging the temperature of response from all 20 cooling cycles. Pearson correlation coefficients were calculated to establish the relationship between the peak-to-peak amplitude and DHEQ score, VAS score, and adjusted temperature of response.

## Results

### Analysis of sensitivity scores and DHEQ scores

All 60 participants (30 in the control group and 30 in the experimental group) were included in the analysis of sensitivity scores and DHEQ scores (Table 1). The DHEQ, which assesses the impact of dentine hypersensitivity on quality of life, showed a strong positive correlation with both sensitivity scores. Specifically, a strong positive correlation was found between the DHEQ score and the clinician-assessed Schiff Sensitivity Scale score for evaporative stimulation (Spearman’s r_s_=0.73, p<0.0001; Figure 3A). Similarly, the DHEQ score strongly correlated with the self-reported VAS score for thermal stimulation (Spearman’s r_s_=0.65, p<0.0001; Figure 3B). To evaluate the relationship between the two different measures of dentine hypersensitivity, a correlation analysis was performed between the clinician-assessed Schiff sensitivity scale score for evaporative stimulation and the self-reported VAS score for thermal stimulation (Figure 3C). This analysis revealed a strong positive correlation between the two sensitivity scores (rs=0.56, p<0.0001; Spearman’s rank correlation), indicating that stronger observed reactions to evaporative stimulation closely align with a higher self-reported intensity of sensation felt during thermal stimulation.

**Table 1.** Clinical information and stimulation response data for all participants (n=60).

| Participant | Group | Age | Gender | Test tooth | Evaporative stimulation |  | Thermal stimulation |  |  | DHEQ score |
| --- | --- | --- | --- | --- | --- | --- | --- | --- | --- | --- |
|  |  |  |  |  | Sensatio n/Pain | Schiff Scale score | Sensatio n/Pain | VAS score | No. cooling cycles with response |  |
| 012 | Control | 55 | M | UR1 | No | 0 | Yes | 1 | 20/20 | 74 |
| 057 | Control | 42 | M | UR3 | No | 0 | Yes | 0.2 | 20/20 | 38 |
| 017 | Control | 64 | M | LR3 | No | 0 | No | 0 | 0/20 | 37 |
| 059 | Control | 23 | F | UR3 | No | 0 | No | 0 | 0/20 | 110 |
| 022 | Experimental | 28 | M | LR5 | Yes | 1 | Yes | 1.2 | 0/20 | 65 |
| 030 | Experimental | 31 | M | UR4 | Yes | 1 | Yes | 4.1 | 8/20 | 120 |
| 060 | Control | 19 | M | UR4 | No | 0 | Yes | 6 | 17/20 | 78 |
| 054 | Experimental | 25 | M | LR1 | No | 0 | Yes | 3 | 19/20 | 75 |
| 023 | Control | 61 | M | UR3 | No | 0 | No | 0 | 0/20 | 50 |
| 062 | Experimental | 44 | F | LR1 | Yes | 1 | Yes | 7 | 19/20 | 181 |
| 041 | Experimental | 23 | M | LL1 | Yes | 1 | Yes | 5.7 | 19/20 | 136 |
| 036 | Control | 61 | M | LR1 | No | 0 | Yes | 0.4 | 1/20 | 37 |
| 052 | Control | 46 | F | UR4 | No | 0 | Yes | 0.4 | 0/20 | 49 |
| 063 | Control | 27 | F | LL1 | No | 0 | Yes | 0.8 | 0/20 | 38 |
| 007 | Experimental | 23 | F | UR4 | Yes | 2 | Yes | 1.5 | 14/20 | 130 |
| 064 | Control | 24 | M | LR5 | No | 0 | Yes | 0.7 | 0/20 | 45 |
| 066 | Control | 21 | F | LR1 | No | 0 | Yes | 1.6 | 0/20 | 36 |
| 068 | Experimental | 21 | F | LR2 | Yes | 1 | Yes | 7.3 | 17/20 | 120 |
| 069 | Control | 22 | F | LR2 | No | 0 | Yes | 3.9 | 7/20 | 37 |
| 008 | Control | 23 | F | LR4 | No | 0 | Yes | 1.1 | 0/20 | 87 |
| 071 | Experimental | 63 | F | LR4 | Yes | 2 | Yes | 4 | 20/20 | 126 |
| 070 | Control | 20 | M | LL4 | No | 0 | Yes | 1.5 | 0/20 | 87 |
| 034 | Experimental | 22 | M | UL5 | Yes | 1 | Yes | 0.6 | 0/20 | 53 |
| 072 | Control | 38 | F | UL5 | No | 0 | Yes | 0.8 | 0/20 | 44 |
| 073 | Experimental | 22 | M | LR1 | Yes | 1 | Yes | 7 | 20/20 | 99 |
| 074 | Experimental | 47 | F | LR5 | Yes | 1 | Yes | 0.8 | 0/20 | 53 |
| 075 | Experimental | 36 | F | UL5 | Yes | 2 | Yes | 6.4 | 18/20 | 161 |
| 076 | Experimental | 18 | F | LL4 | Yes | 1 | Yes | 3.3 | 0/20 | 80 |
| 077 | Control | 23 | F | LR1 | No | 0 | Yes | 0.7 | 0/20 | 36 |
| 078 | Experimental | 30 | F | LL1 | Yes | 1 | Yes | 4.1 | 0/20 | 93 |
| 055 | Control | 22 | M | LL1 | No | 0 | Yes | 3.9 | 0/20 | 51 |
| 079 | Control | 26 | F | LR5 | No | 0 | Yes | 5 | 0/20 | 58 |
| 080 | Experimental | 36 | F | LL5 | Yes | 2 | Yes | 1.7 | 0/20 | 161 |
| 081 | Experimental | 25 | M | LL4 | Yes | 1 | Yes | 1.1 | 0/20 | 59 |
| 082 | Experimental | 19 | F | LR1 | Yes | 1 | Yes | 5.5 | 20/20 | 113 |
| 083 | Control | 63 | F | LL5 | No | 0 | No | 0 | 0/20 | 38 |
| 084 | Experimental | 51 | M | UR5 | Yes | 1 | Yes | 1.3 | 0/20 | 85 |
| 085 | Experimental | 48 | F | LR5 | Yes | 2 | Yes | 7.4 | 19/20 | 161 |
| 086 | Experimental | 54 | M | UR5 | Yes | 2 | Yes | 5.3 | 0/20 | 122 |
| 088 | Control | 18 | M | LL4 | No | 0 | Yes | 5.7 | 1/20 | 68 |
| 089 | Experimental | 44 | M | LR1 | Yes | 1 | Yes | 5.5 | 20/20 | 129 |
| 090 | Control | 30 | F | LR1 | No | 0 | Yes | 3.8 | 17/20 | 143 |
| 091 | Control | 20 | F | UL5 | No | 0 | Yes | 1.9 | 0/20 | 47 |
| 003 | Experimental | 23 | M | UR5 | Yes | 1 | Yes | 5.7 | 2/20 | 148 |
| 092 | Control | 43 | F | UR5 | No | 0 | Yes | 1.4 | 17/20 | 39 |
| 093 | Control | 25 | F | UR5 | No | 0 | Yes | 2.1 | 1/20 | 102 |
| 094 | Experimental | 61 | F | LR1 | Yes | 1 | Yes | 5.1 | 20/20 | 115 |
| 095 | Experimental | 53 | F | LR2 | Yes | 2 | Yes | 8.8 | 8/20 | 58 |
| 096 | Experimental | 57 | F | UL4 | Yes | 2 | Yes | 7 | 20/20 | 194 |
| 097 | Control | 21 | M | UL4 | No | 0 | Yes | 4.7 | 12/20 | 39 |
| 087 | Control | 28 | F | UR5 | No | 0 | Yes | 2.4 | 0/20 | 96 |
| 098 | Control | 24 | F | LR5 | No | 0 | Yes | 4.1 | 9/20 | 37 |
| 099 | Experimental | 20 | F | UR1 | Yes | 2 | Yes | 6.9 | 20/20 | 158 |
| 100 | Experimental | 52 | M | UR5 | Yes | 2 | Yes | 3.8 | 12/20 | 98 |
| 101 | Experimental | 46 | F | LL3 | Yes | 2 | Yes | 4.5 | 19/20 | 190 |
| 010 | Experimental | 53 | F | UR2 | Yes | 2 | Yes | 6.9 | 20/20 | 130 |
| 102 | Control | 22 | M | LR1 | No | 0 | Yes | 5.1 | 18/20 | 57 |
| 020 | Experimental | 58 | M | UL5 | Yes | 2 | Yes | 5.1 | 20/20 | 109 |
| 002 | Control | 42 | M | UL5 | No | 0 | No | 0 | 0/20 | 35 |
| 103 | Control | 50 | F | LL3 | No | 0 | Yes | 0.2 | 0/20 | 37 |

**Figure 3.**
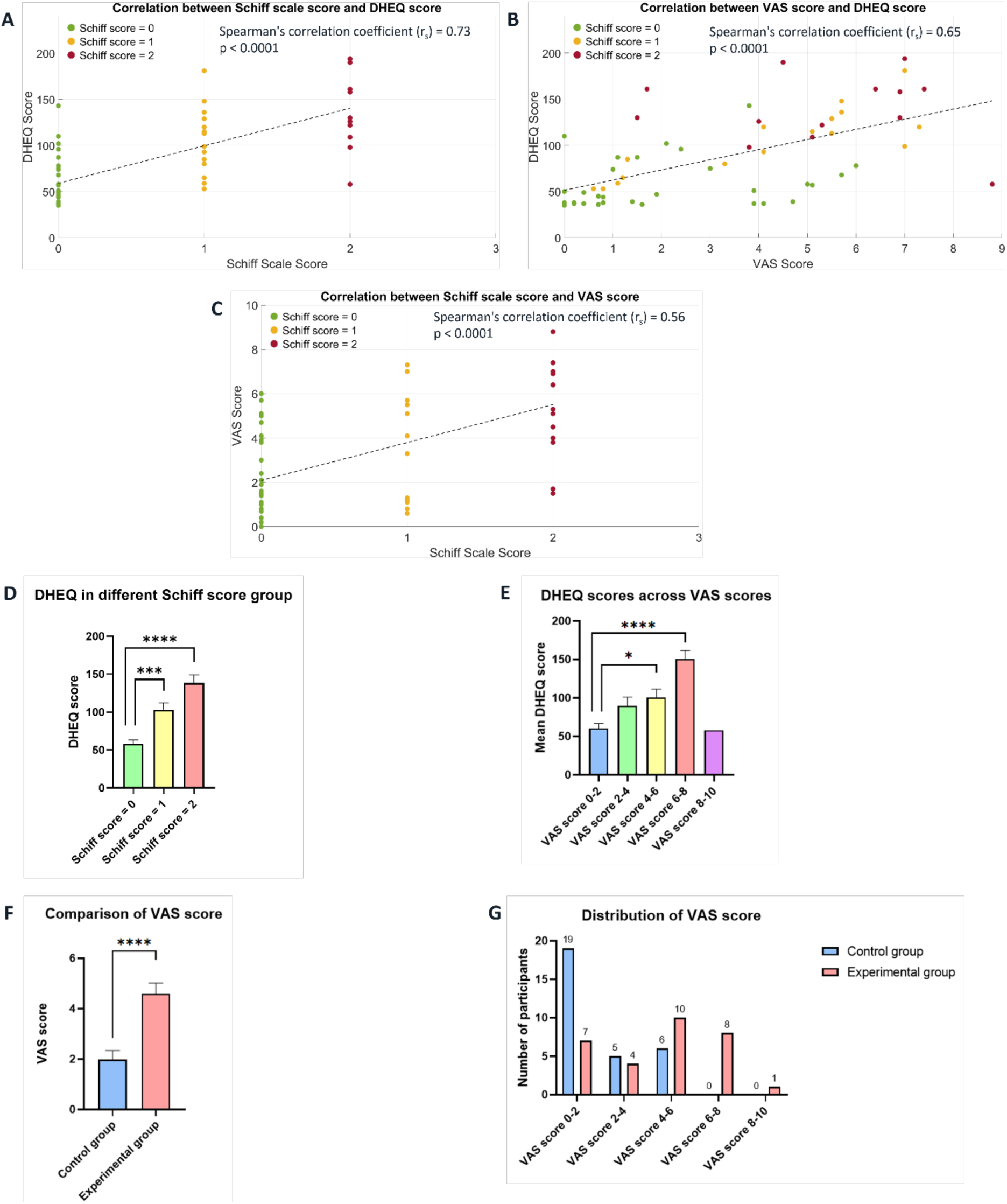
Correlation and group differences for the DHEQ and sensitivity scores. A) Following evaporative stimulation, the clinician assessed the participant’s response to the air stimuli using the Schiff Sensitivity Scale. There is a strong positive correlation between the DHEQ score and the clinician-assessed Schiff sensitivity scale score for evaporative stimulation (rs=0.73, p<0.0001; Spearman’s rank correlation). B) Following thermal stimulation, participants rated the intensity of sensation they felt by completing the VAS scale. There is a strong positive correlation between the DHEQ score and the self-reported VAS score for thermal stimulation (rs=0.65, p<0.0001; Spearman’s rank correlation). The Schiff Scale scores for each participant were indicated by different colours: Green = Schiff score = 0; Yellow = Schiff score = 1; Red = Schiff score = 2. C) Scatter plot illustrating a strong positive correlation between the clinician-assessed Schiff sensitivity scale score for evaporative stimulation and the self-reported VAS score for thermal stimulation (rs=0.56, p<0.0001; Spearman’s rank correlation). The Schiff Scale scores for each participant were indicated by different colours: Green = Schiff score = 0; Yellow = Schiff score = 1; Red = Schiff score = 2. D) Comparison of DHEQ scores (mean ± SEM) across participants with different Schiff scores (0, 1, and 2). Kruskal-Wallis ANOVA followed by Dunn’s multiple comparisons test showed that participants with Schiff score of 1 and Schiff score of 2 had significantly higher DHEQ scores compared to participants with Schiff score of 0 (p<0.001 and p<0.0001, respectively). E) Comparison of DHEQ scores (mean ± SEM) across participants with different VAS score categories (0 – 2, 2 – 4, 4 – 6, 6 – 8, and 8 – 10). Kruskal-Wallis ANOVA followed by Dunn’s multiple comparisons test showed that participants with VAS scores of 4 – 6 and 6 – 8 had significantly higher DHEQ scores compared to participants with VAS score of 0 – 2 (p=0.026 and p<0.0001, respectively). F) Comparison of VAS scores (mean ± SEM) between control (no dentine hypersensitivity) and experimental group (with dentine hypersensitivity). Participants in the experimental group reported significantly higher VAS score than participants in the control group (p<0.0001; unpaired t-test). G) Distribution of VAS scores reported by participants in the control and experimental group, showing a clear shift towards higher VAS scores reported by the experimental group.

To further evaluate the relationship between DHEQ score and sensitivity scores, DHEQ scores were compared across the three Schiff score groups (Figure 3D) and different VAS score categories (Figure 3E). Participants with Schiff scores of 1 and 2 had significantly higher DHEQ scores compared to those with a Schiff score of 0 (p<0.001 and p<0.0001, respectively; Kruskal-Wallis ANOVA with Dunn’s multiple comparisons test). On the other hand, participants who rated the sensation they felt as more painful (VAS scores of 4–6 and 6–8) had significantly higher DHEQ scores than those who reported less or minimal discomfort (VAS scores of 0–2) (p=0.026 and p<0.0001, respectively; Kruskal-Wallis ANOVA with Dunn’s multiple comparisons test).

Subjective discomfort associated with thermal stimulation was assessed using the VAS score upon completion of thermal sensitivity assessment. Participants with dentine hypersensitivity (experimental group) reported significantly higher VAS scores compared to participants with no dentine hypersensitivity (control group) (p<0.0001; unpaired t-test; Figure 3F), which suggests that they felt greater discomfort. This is also shown in the distribution of VAS scores reported by both groups of participants (Figure 3G), which demonstrated a shift towards higher discomfort ratings in the experimental group.

### Comparison of EEG responses to evaporative stimulation by Schiff scores

Average event-related potential (ERP) traces were calculated for 59 participants (data from one participant was excluded from the analysis due to poor signal quality, as detailed in the methods) and were compared based on the severity of the response to air puffs (Schiff scores: 0, n=30; 1, n=16; and 2, n=13; Figure 4). The time course of the ERPs was analysed using Welch’s t-test on sequential 100 ms windows (overlapping by 50 ms), corrected for multiple comparisons (α=0.0167).

**Figure 4.**
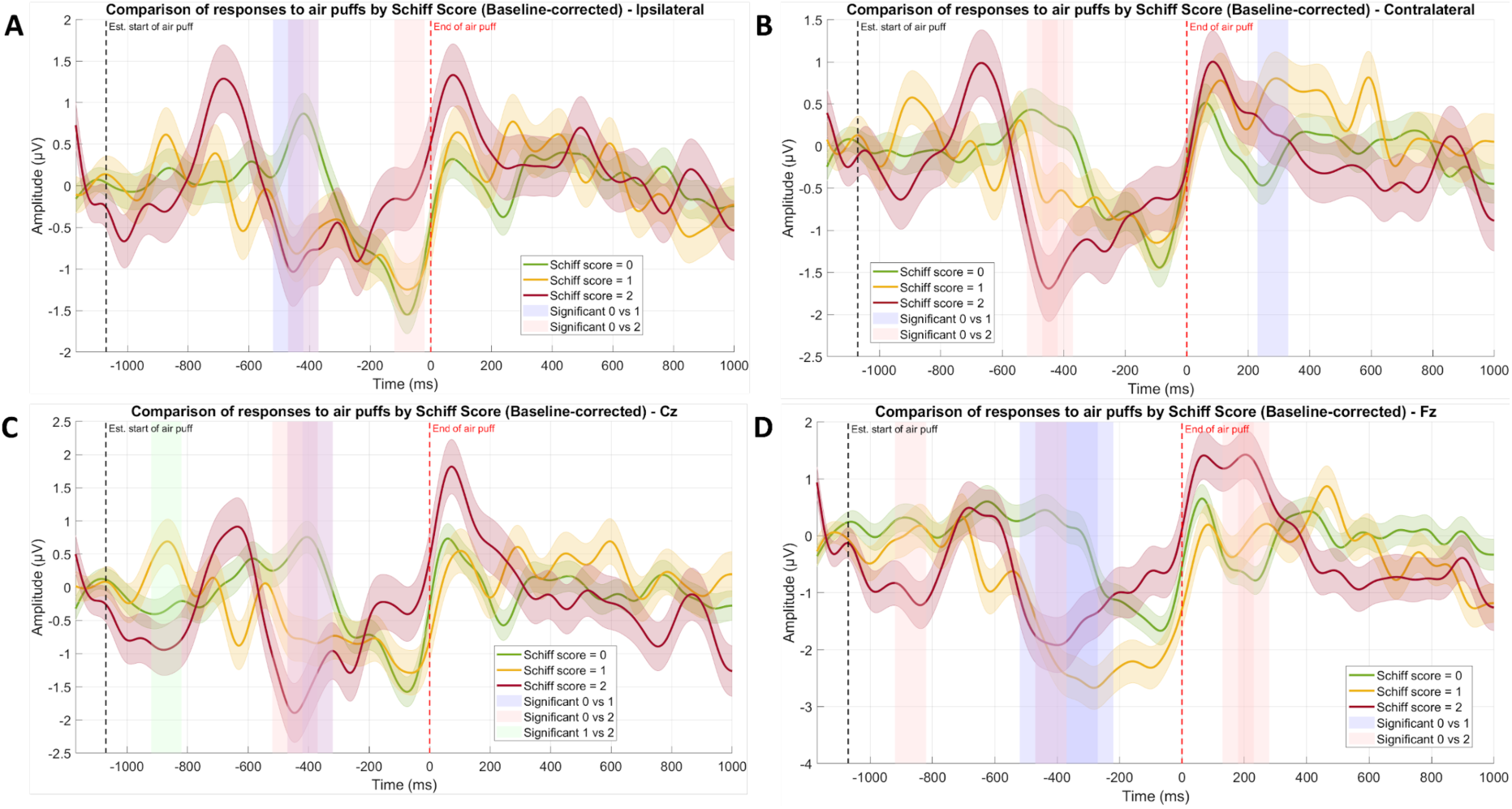
Comparison of event-related potential (ERP) responses to evaporative stimulation across Schiff score groups. Average ERP traces (mean ± SEM) for all participants that were included in data analysis (n=59) in response to evaporative stimulation (air puff) across participants with different Schiff scores (Schiff score = 0, n=30; Schiff score = 1, n=16; Schiff score = 2, n=13). ERPs were baseline-corrected relative to the 100 ms window immediately preceding the estimated air puff onset (black dashed line at - 1070 ms). The trials were time-locked to the air puff offset (red dashed line at 0 ms). Data are shown for the four channels used in the analysis: (A) Ipsilateral somatosensory cortex (C3/C4 depending on the side of the test tooth), (B) Contralateral somatosensory cortex (C3/C4 depending on the side of the test tooth), (C) Cz, (D) Fz. Statistical significance was assessed using Welch’s t-test on sequential 100 ms time windows (overlapping by 50 ms), corrected for multiple comparisons (α=0.0167, Bonferroni corrected for 3 pairwise comparisons). Shaded time windows indicate significant difference in mean amplitude between Schiff score groups: Blue = Schiff 0 vs Schiff 1; Red = Schiff 0 vs Schiff 2; Green = Schiff 1 vs Schiff 2.

The analysis revealed distinct ERP profiles across the Schiff score groups, with significant differences mainly concentrated in the period when air puffs were applied (-1070 ms to 0 ms relative to air puff offset). In particular, the most pronounced and widespread differences across all four analysed channels occurred during the period between -470 ms and -370 ms relative to air puff offset (Figure 4). This period is characterised by a significant difference in mean amplitude, where the sensitive groups (Schiff scores of 1 and 2) consistently exhibited a significantly more negative deflection compared to the non-sensitive group (Schiff score of 0), reflecting enhanced cortical activity in participants with dentine hypersensitivity.

Furthermore, late positivity was observed after the cessation of the evaporative stimulus. In the contralateral somatosensory cortex, the ERP from participants with Schiff score of 1 exhibited a significantly more positive deflection compared to that from participants with Schiff score of 0 in the period from +230 ms to +330 ms relative to air puff offset (Figure 4B). Similarly, the ERP in Fz from participants with Schiff score of 2 exhibited a significantly more positive deflection compared to that from participants with Schiff score of 0 in the period from +130 ms to +280 ms relative to air puff offset (Figure 4D). These indicate the presence of a post-stimulus cortical response following stimulus offset.

### EEG responses to thermal stimulation

#### Temporal dynamics of response and no response trials

EEG data from the 59 participants were separated into two groups based on trial outcome, resulting in a total of 324 valid response trials (where participants pressed a button to indicate discomfort) and 541 valid no response trials (where the cooling cycle finished without a response). To isolate processing associated with nociception and motor execution of button press, cortical activity associated with the button press was subtracted from the response trials prior to analysis.

The average ERP traces for the two trial outcomes were compared in Figure 5, showing the 4-second window spanning from -3000 ms to +1000 ms relative to the trial termination marker. For all four analysed channels (Figure 5), the no response trials exhibited a relatively stable trace throughout this time window. In contrast, the response trials displayed a distinct change in cortical activity, primarily characterised by a slow, sustained negative potential followed by a large-amplitude positive deflection (as detailed below), which is primarily observed in the contralateral somatosensory cortex and Fz. Statistical analysis using Welch’s t-test on sequential 100 ms windows (overlapping by 50 ms) revealed significant differences between response and no response trials.

**Figure 5.**
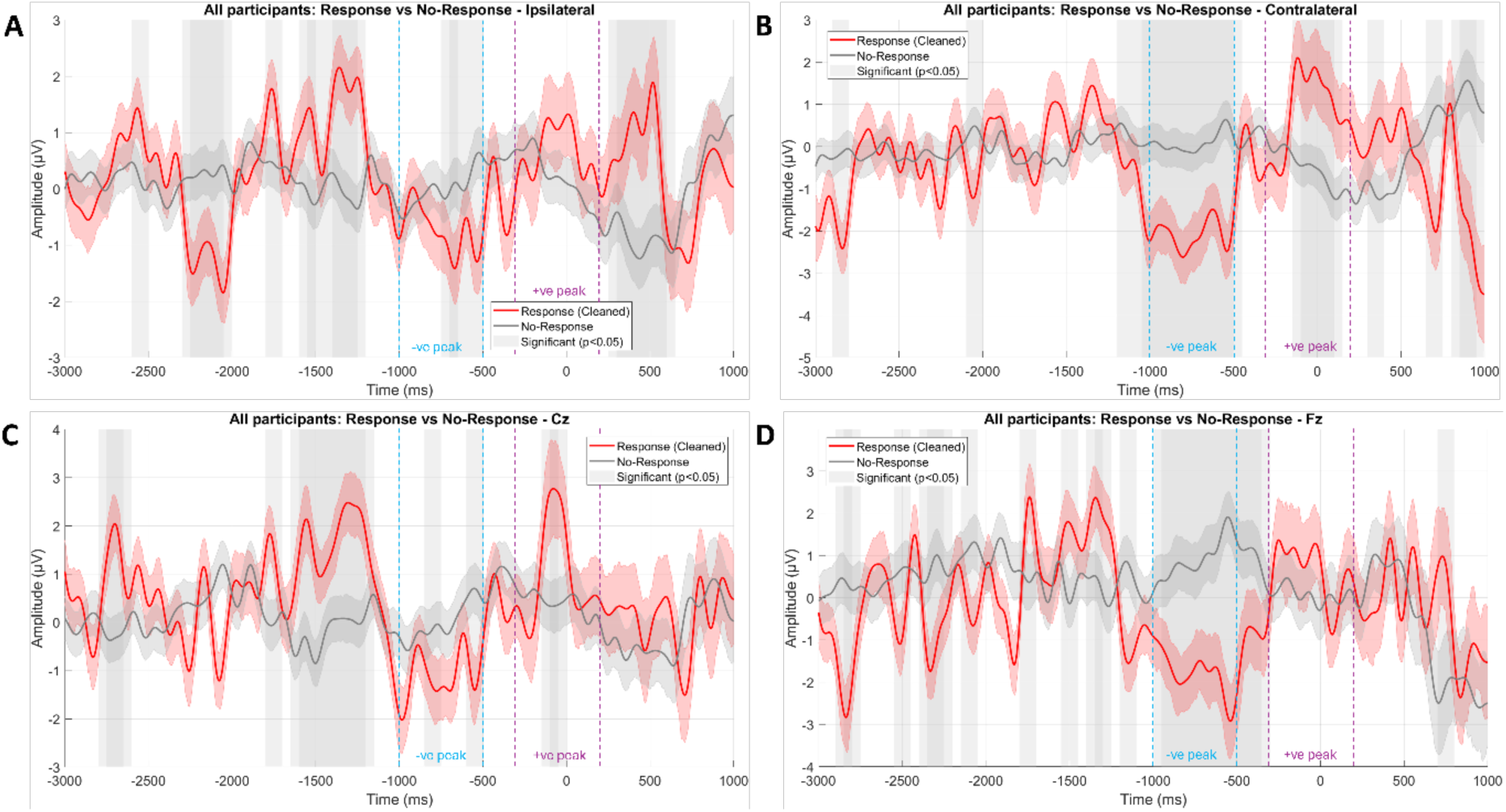
Comparison of ERP responses to thermal stimulation between response and no response trials. Average ERP traces (mean ± SEM) for all participants that were included in data analysis (n=59) during cooling cycles. Traces were separated by trial outcome: Response (cooling cycles where participants pressed ‘s’ to indicate discomfort; n=324) and No response (cooling cycles completed without a response; n=541). All traces were baseline-corrected relative to the 2-second baseline window that began 1 second after the trial’s start marker. For response trials, cortical activity associated with the motor execution of button press was subtracted prior to analysis in order to isolate processing associated with nociception. Trials were time-locked to the trial termination marker (at 0 ms), which corresponds to either the participant’s button press (Response) or end of the cooling cycle (No response). Data are shown for the four channels used in the analysis: (A) Ipsilateral somatosensory cortex (C3/C4 depending on the side of the test tooth), (B) Contralateral somatosensory cortex (C3/C4 depending on the side of the test tooth), (C) Cz, (D) Fz. Statistical significance was assessed using Welch’s t-test on sequential 100 ms time windows (overlapping by 50 ms), with a significance threshold of p<0.05. Grey shading indicates time windows with a statistically significant difference in mean amplitude between the response and no response groups. Blue and purple dotted lines mark how the minimum and maximum amplitude around the trial termination marker were obtained.

The 3000 ms pre-termination window relative to the trial termination marker captured the sustained neural activity prior to the immediate decision/reaction phase. During this period, there are significant deviations between response and no response trials, though the morphology varied by region. EEG waveforms from the contralateral somatosensory cortex (Figure 5B) and Fz (Figure 5D) exhibited a consolidated period of sustained significant negative potential in the response trials. In the contralateral somatosensory cortex, this negativity in the response trials spanned from -1100 ms to -450 ms relative to the trial termination marker. Similarly, the ERPs at Fz exhibited a significant negative period in the response trials that spanned from -1000 ms to -300 ms. In contrast, EEG waveforms from the ipsilateral somatosensory cortex (Figure 5A) and Cz (Figure 5C) displayed a more complex pattern in the response trials. ERPs from the ipsilateral somatosensory cortex exhibited a distinct alternating profile in the response trials: a sustained significant negative potential (-2300 ms to -2000 ms) was followed by a significant positive deflection (-1600 ms to -1200 ms), before returning to a brief negative deflection (-750 ms to -600 ms) prior to the response (at 0 ms). In Cz, the response trials showed an initial period of significant positive deflection (-2800 ms to - 2600 ms) followed by a prolonged period of negativity (-1650 ms to -1150 ms).

The 1000 ms post-termination window relative to the trial termination marker captured the neural activity associated with the response and immediate aftermath of the decision. During this period, the response trials were characterised by a positive deflection across all channels, though the statistical significance of the difference relative to no response trials varied by region. The ipsilateral somatosensory cortex (Figure 5A) exhibited a significant positive period in the response trials, spanning from +250 ms to +650 ms. In the contralateral somatosensory cortex (Figure 5B), significant positive deflections were observed in the response trials relative to no response trials in an earlier window immediately preceding the response that spanned from -150 ms to +150 ms, followed by a brief significant positive deflection that spanned from +300 ms to +400 ms. Similarly in Cz (Figure 5C), a significant positive deflection in the response trials immediately preceding the response from -150 ms to 0 ms was observed. Notably, although the ERP waveform at Fz showed a visually distinct positive deflection in the response trials at around 0 ms, this component was not statistically significantly different from the no response trials.

#### Topographical analysis of EEG band power

Topographical maps of EEG band power depict the spatial distribution and intensity of neural oscillations across the scalp. To visualise the spatial distribution and temporal evolution of cortical activation in the period preceding the participants’ decision to respond, topographical maps of mean EEG band power at every 0.5-second time window were generated for the 5-second period preceding trial termination for both response and no response trials (Figure 6).

**Figure 6.**
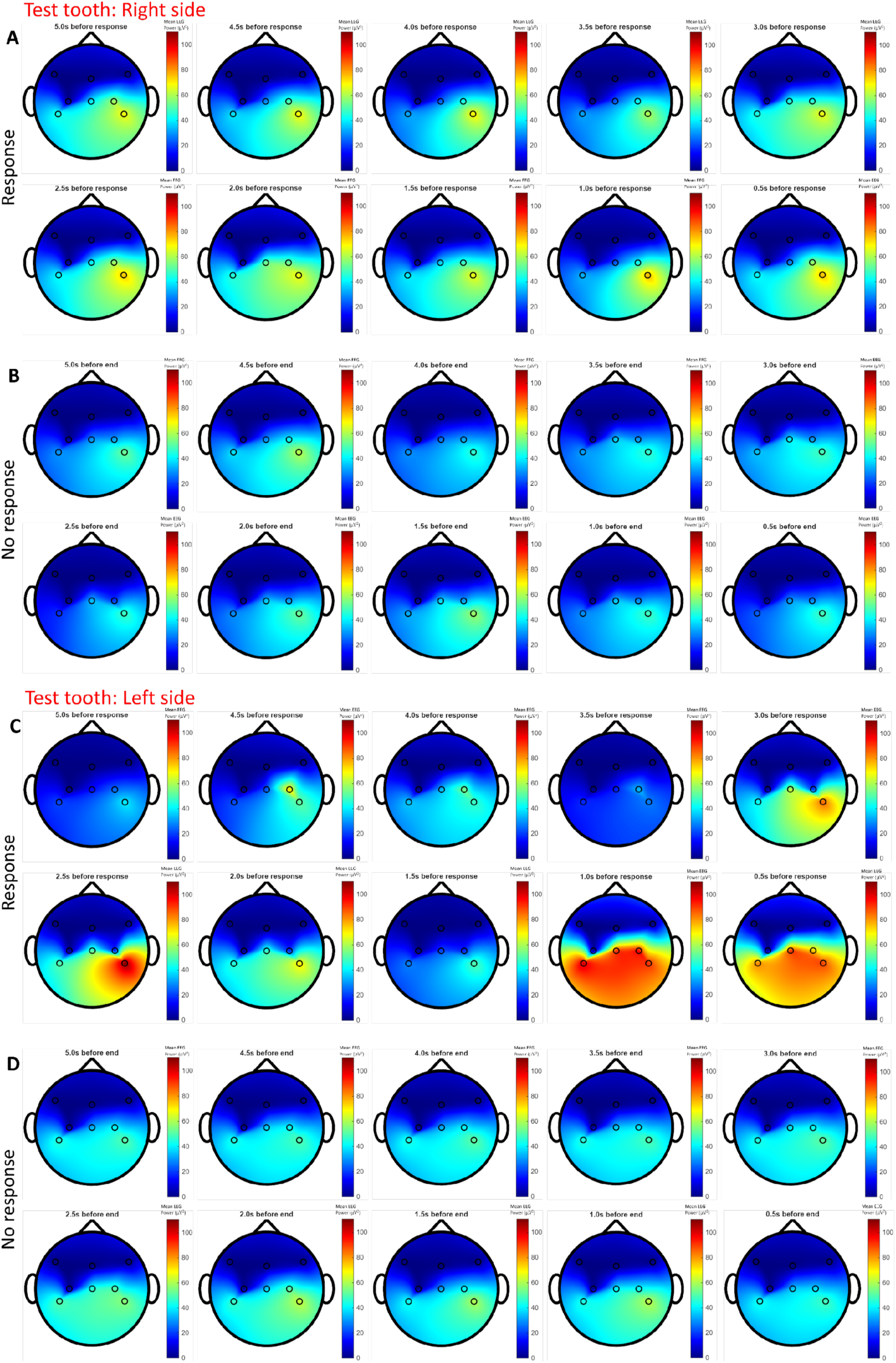
Topographical distribution of EEG band power prior to trial termination for response and no response trials. Topographic maps illustrating the spatial distribution of mean EEG band power across the scalp in the 5-second period preceding trial termination. Maps were generated after baseline correction and, for response trials, point-by-point subtraction of motor execution of button press. Data were averaged across participants (Right side: n=39; Left side: n=20) and are presented in ten consecutive 0.5 s time segments (from -5 s to -0.5 s relative to trial termination). The maps were grouped by trial outcome (Response: A, C; No response: B, D) and the side of the test tooth (Right side: A-B; Left side: C-D). Response trials where the test tooth was on the right (A) and left (C) side both showed a progressive increase in EEG band power leading up to the response, whereas no response trials for the right (B) and left (D) side both showed a relatively stable, low-power topography. The colour scale represents the band power (µV^2^), with warmer colours (red, yellow) indicating higher power, while cooler colours (blue, green) indicate lower power.

For trials where the test tooth was on the right side (Figures 6A–B), the response trials (Figure 6A) exhibited a progressive increase in band power that was predominantly lateralised to the ipsilateral (right) central parietal region (CP6), an area associated with integrating sensorimotor information [11]. The EEG power became increasingly focal in the final 1 second leading up to the response. This contrasts with the no response trials (Figure 6B), which maintained a relatively stable, low-power topography with no distinct focal activation throughout the 5-second period.

For trials where the test tooth was on the left side (Figures 6C-D), the response trials (Figure 6C) also displayed a progressive increase in power that was initiated in the contralateral (right) central parietal region (CP6) 3 seconds prior to the response. This activation subsequently spread to involve the ipsilateral (left) central parietal region (CP5), resulting in a bilateral recruitment that intensified in the final 1 second leading up to the response. This contrasts with the spatially diffuse and relatively stable topography observed in the no response trials (Figure 6D).

Across response trials from both sides (Figure 6A, 6C), a progressive pattern of cortical recruitment that varied by laterality was observed, yet both followed a clear temporal gradient. The earliest sign of focal activation emerged 3 seconds prior to the response, which intensified in the final 1 second window leading up to the response. This spatiotemporal profile suggests that the decision to signal discomfort is accompanied by the focal onset and progressive expansion of cortical excitability originating from areas associated with sensory processing [11].

#### Comparison of EEG band power between response and no response trials

EEG waveforms are categorised by their frequencies into five different frequency bands: delta (1–4 Hz), theta (4–8 Hz), alpha (8–12 Hz), beta (12–30 Hz), and gamma (30–100 Hz). To identify the specific frequency bands underlying the cortical activity associated with the participants’ decision to indicate discomfort, the mean band power was quantified, which reflects the intensity of neural oscillations within specific frequency bands. Mean EEG band power was compared between response and no response trials for the 2-second window immediately preceding trial termination (Figure 7). This analysis showed that there was a frequency-specific modulation of neural oscillations associated with the response.

**Figure 7.**
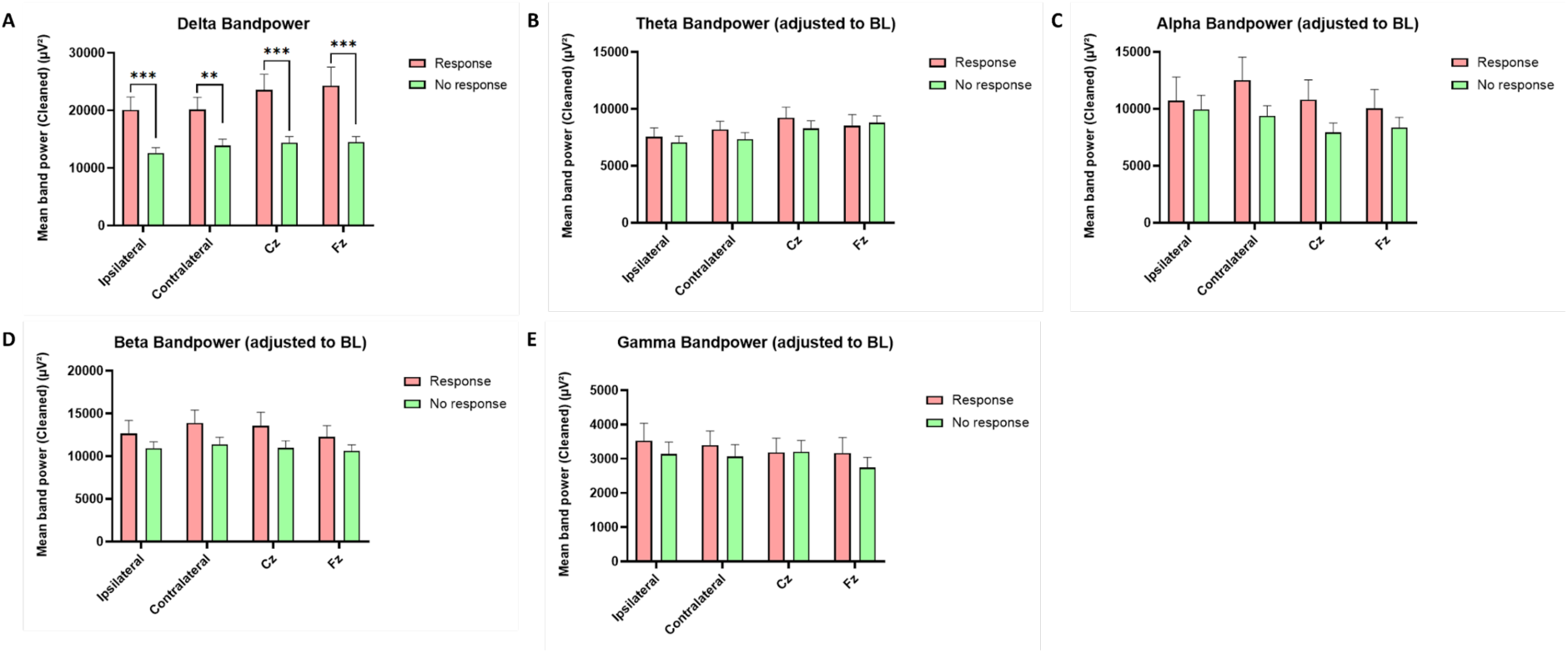
Comparison of EEG band power between response and no response trials. Comparison of mean band power for response (red) and no response (green) trials across five frequency bands: delta (1–4 Hz; A), theta (4–8 Hz; B), alpha (8–12 Hz; C), beta (12–30 Hz; D), and gamma (30–100 Hz; E). Data were averaged across all participants (n=59) and calculated for the 2-second time window immediately preceding the trial termination marker, following baseline correction and, for response trials, point-by-point subtraction of motor execution of button press. Statistical comparisons were assessed using unpaired t-tests or Mann-Whitney tests, depending on the normality of data. A statistically significant increase in power was observed only in the delta band (A) for response trials in comparison to no response trials across all four channels (**p<0.01, ***p<0.001). No significant differences were found for the theta, alpha, beta, or gamma bands.

In the delta frequency band (Figure 7A), the mean band power in response trials was significantly higher compared to no response trials across all four analysed channels. This was revealed by unpaired t-tests in the ipsilateral somatosensory cortex (p<0.001) and Cz (p<0.001), and by Mann-Whitney tests in the contralateral somatosensory cortex (p=0.008) and Fz (p=0.001). In contrast, no statistically significant differences between response and no response trials were observed for the theta (Figure 7B), alpha (Figure 7C), beta (Figure 7D), and gamma (Figure 7E) frequency bands in any of the four channels (Figures 7B-E). These findings suggest that the cortical processing of the cold stimulus and the subsequent decision to signal discomfort are primarily characterised by a widespread elevation in lower-frequency delta activity, rather than modulation in higher frequency bands.

#### Correlation between peak-to-peak amplitude and measures of sensitivity

Peak-to-peak amplitude was defined as the difference between the minimum and maximum amplitude around the trial termination marker (minimum: -1000 to -500 ms; maximum: -300 to +200 ms), as indicated on Figure 5. In a previous study, it was found that peak-to-peak amplitude had a significant positive correlation with the participants’ estimate of thermal stimulus intensity applied to the back of their hand [12]. To investigate whether the peak-to-peak amplitude serves as a suitable objective marker for dentine hypersensitivity, Pearson correlation analyses were performed between the peak-to-peak amplitudes and measures of sensitivity (DHEQ, VAS score, and temperature of response).

At Fz, the peak-to-peak amplitude exhibited a significant positive correlation with both the DHEQ score (r=0.3, p=0.04; Figure 8A) and the VAS score (r=0.29, p=0.048; Figure 8B). This suggests that a larger magnitude of the cortical response in the central frontal region is associated with greater impact of dentine hypersensitivity on quality of life and greater intensity of sensation felt by participants. However, no significant correlations were observed for the DHEQ (Figure 8A) or VAS (Figure 8B) scores in the ipsilateral somatosensory cortex, contralateral somatosensory cortex, or Cz.

**Figure 8.**
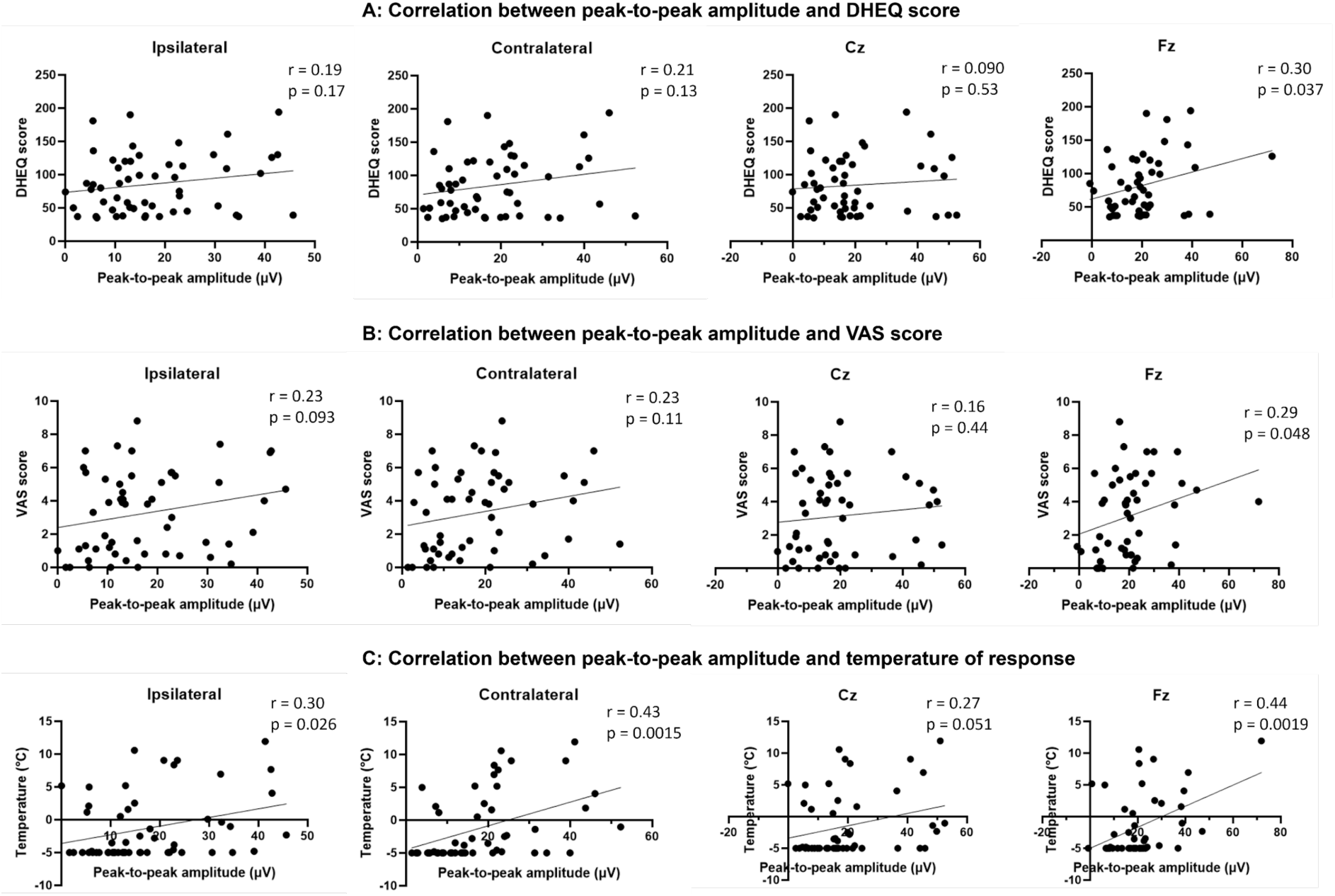
Correlations between peak-to-peak amplitude and sensitivity measures. Scatter plots illustrating the relationship between the peak-to-peak amplitude of the ERP response to thermal stimulation and the DHEQ score (A), VAS score (B), and temperature of response (C). Data points represent individual participants (n=59). Peak-to-peak amplitude was calculated as the difference between the maximum positive peak (-300 ms to +200 ms relative to trial termination marker) and the minimum negative peak (-1000 ms to -500 ms relative to trial termination marker), as indicated on Figure 5. Relationships were assessed using Pearson correlation coefficient (r) for each of the four channels: Ipsilateral, Contralateral, Cz, and Fz. (A) Correlation between peak-to-peak amplitude and the DHEQ score, a measure of the overall impact of dentine hypersensitivity on quality of life. (B) Correlation between peak-to-peak amplitude and the self-reported VAS score. (C) Correlation between peak-to-peak amplitude and the adjusted temperature of response (averaged across all trials, with no response trials set to -5°C). Pearson correlation coefficients (r) and significance levels (p) are displayed within each panel.

A more widespread relationship with peak-to-peak amplitude was observed for the adjusted temperature of response (Figure 8C). Significant positive correlations were found between the peak-to-peak amplitude and the adjusted temperature of response in the ipsilateral somatosensory cortex (r=0.30, p=0.03), contralateral somatosensory cortex (r=0.43, p=0.002), and Fz (r=0.44, p=0.002). This positive correlation indicates that participants who responded to thermal stimulation at higher temperatures (lower threshold of thermal sensitivity) exhibited significantly larger cortical responses in the somatosensory and frontal regions compared to those who responded at lower temperatures (higher threshold of thermal sensitivity). No significant correlation was observed for the adjusted temperature of response in Cz.

## Discussion

This study aimed to determine whether EEG responses to tooth stimulation can serve as an objective measure of dentine hypersensitivity severity, and whether subjective ratings of thermal stimulation (VAS score) can effectively assess the condition. Our findings revealed that both the clinician-assessed Schiff sensitivity score and the self-reported VAS score had a strong positive correlation with the DHEQ score. Notably, participants who had a stronger response to air puffs (higher Schiff scores as assessed by the clinician), as well as those who reported the intensity of sensation felt during thermal stimulation as more painful, had significantly higher DHEQ scores than participants who did not respond to air puffs and those who reported minimal discomfort during thermal stimulation, respectively. These findings demonstrate that both the Schiff score and the VAS score can accurately capture the impact of dentine hypersensitivity.

Moreover, the results demonstrated that EEG responses to evaporative stimulation (air puffs) differed significantly between participants with different Schiff scores. Similarly, EEG responses to thermal stimulation differed significantly between trials where participants indicated discomfort and those where they did not. Taken together, these findings suggest an association between cortical activity and measures of dentine hypersensitivity.

### Cortical responses to evaporative stimulation

Air puffs applied to sensitive teeth (Schiff score ≥ 1) evoked changes in ERP responses that were significantly different from those in non-sensitive teeth (Schiff score = 0) in the somatosensory cortex (both ipsilateral and contralateral to the test tooth), the primary motor cortex (Cz), and the central frontal region (Fz) of the brain. Significant differences in brain activity, recorded using functional magnetic resonance imaging (fMRI), were also observed when sensitive teeth were stimulated with air stimuli in comparison to when non-sensitive teeth were stimulated [13]. In addition to spatial activation in response to air puffs[13], our study showed the temporal dynamics of this differential processing. Specifically, both sensitive groups (Schiff score of 1 and 2) exhibited a more pronounced negativity than the non-sensitive group at approximately 400 ms prior to stimulus offset across the somatosensory, motor, and frontal regions, which may reflect the enhanced nociceptive processing and discomfort experienced by participants with sensitive teeth when air puffs were applied. This aligns with findings in a previous study that investigated laser-evoked EEG responses to noxious heat stimuli, where a strong positive correlation was found between the magnitude of negative potentials elicited by the stimulus and the intensity of perceived pain rated by the participants [14]. This increased negativity in the sensitive groups may therefore be indicative of greater discomfort induced by air puffs.

In addition to the sustained negativity during air puff application, a significant late positivity following stimulus termination was identified in the central frontal region (Fz) in participants with a Schiff score of 2, who reacted significantly to air puffs. This post-stimulus positivity might reflect an attentional shift from the discomfort brought by the air puff to processing the cessation of the auditory signals when the hissing sound associated with the air puff stopped [15]. This is supported by a previous study on the P3a component of the P300, a frontal positive potential occurring 250–500 ms after stimulus offset that reflects the shift of attention to novel or changing sensory events [16]. The presence of this late positive component in participants with a Schiff score of 2 therefore suggests that these participants were more aware of the termination of the evaporative stimuli that are noxious to them in comparison to other participants. Taken together, these findings suggest that air puff stimuli elicit a multi-component cortical response where the earlier negativity indicates the intensity of sensation, while the later positivity in the frontal region likely reflects the detection of stimulus termination. In future studies, responses to the sound of the air puff could be controlled for by including trials with air puff sound, but not accompanied by air puff stimulation of the tooth. This would help to better distinguish the EEG response associated with the discomfort felt by participants to responses related to the noise of the air puff.

It is important to note that the start point of the air puff stimulus was estimated due to a technical error. Specifically, for the first 48 participants, only the end times of the air puffs were recorded, so the start times were calculated backwards using an average duration derived from the remaining participants where both start and end markers were correctly recorded. While this estimation was applied to all participants to ensure that all data could be analysed consistently, it assumes that the duration of every air puff was identical, which does not account for the minor natural variations in air puff duration that arise from the clinician manually timing the 1-second application.

Consequently, the precise alignment of EEG activity to the onset of air puff has limited accuracy. Therefore, components of ERP including the N2 and P2 (the second negative and positive peaks of the EEG response typically observed after a stimulus [17]), which rely on precise timing relative to stimulus onset, could not be reliably calculated or analysed based on the data collected. Thus, correlation analysis between the peak-to-peak amplitude (often derived from the difference between N2 and P2) and the Schiff score could not be performed.

### Cortical responses to thermal stimulation

#### Temporal dynamics of cortical responses

In trials where participants indicated discomfort, the EEG response to thermal stimulation was characterised by distinct pre-response activity. Most notably, a sustained negative potential emerged approximately 1 second before the response in the contralateral somatosensory cortex and central frontal region (Fz), which differed significantly from trials with no response. The negative deflection in the contralateral somatosensory cortex likely reflects the discomfort felt by the participants. This aligns with a previous study that recorded EEG activity during thermal laser stimulation on the back of the participant’s right hand, where it was found that the most negative deflection after the stimulus onset and prior to responding to noxious stimuli was maximal at C3 (contralateral somatosensory cortex to the stimulated hand) [18]. The study also identified a positive correlation between the amplitude of this negative wave and the participants’ reported perceived pain intensity rating, indicating that a larger amplitude corresponded to a higher perceived pain rating [18]. Furthermore, a study utilising magnetoencephalography to record brain activity while applying thermal laser stimuli to the back of the participant’s hand demonstrated that activation in the contralateral primary somatosensory cortex had a significant positive correlation with the intensity of the applied stimulus [19]. This relationship matched closely with the participants’ pain ratings, which indicates that the contralateral somatosensory cortex may also act to evaluate the magnitude of discomfort [19] prior to making the decision to press ‘s’ to indicate discomfort. The sustained negative potential in the central frontal region may also be linked to the anticipation and planning of movement, as findings from another study also found sustained negativity at Fz before participants press a button to indicate the direction of motion of a dot [20], which suggests that the neural activities may represent the formation of the intention to press ‘s’ to terminate the cooling cycle and indicate discomfort.

In addition, EEG response to thermal stimulation was characterised by distinct post-response activity in trials where participants indicated discomfort. Around the time when participants pressed ‘s’ to terminate the cooling cycle, a significant positive wave emerged across the ipsilateral somatosensory cortex, the contralateral somatosensory cortex, and the central frontal region, which differed significantly from trials with no response. Since processing associated with the motor execution of button press was subtracted from the response trials prior to analysis, these late positive waves are unlikely to be caused solely by motor movement. Instead, they likely represent the brain processing the cessation of the noxious nociceptive input and potentially the subsequent relief once the thermal probe was removed from the tooth at 0 ms [16]. However, while efforts were made to isolate cortical activity associated with nociception from that of the motor execution of button press, variations in motor behaviour may still exist. Specifically, the urgent action of pressing ’s’ to terminate the cooling cycle that caused discomfort may differ from the deliberate button presses performed after counting 10 to 20 seconds during the button press control trials. While the variance in motor execution cannot be completely ruled out, the emergence of the positivity around and after the response strongly points to the cognitive evaluation of relief from discomfort. Beyond these temporal dynamics, examining the spatial distribution of EEG band power in the period preceding trial termination provides further insights into how the brain maps the discomfort associated with cold stimulation, discussed below.

#### Spatial distribution of cortical activity

Topographic maps illustrating the spatial distribution of mean EEG band power across the scalp in the 5-second period before participants indicated discomfort revealed distinct patterns of cortical recruitment that varied depending on the side of the mouth where the stimulus was applied. In EEG analysis, an increase in band power reflects enhanced synchronisation of underlying neuronal populations, so it serves as a reliable marker of heightened regional brain activity [21]. Therefore, the regions exhibiting the greatest increase in band power in the seconds leading up to the response represent the neural networks most actively engaged in processing the accumulating discomfort associated with cold stimulation. Since the somatosensory pathways cross the midline, sensory input from one side of the body is typically processed most prominently by the contralateral cerebral hemisphere. Consistent with these anatomical pathways, our results from trials where participants indicated discomfort showed that cold stimulation of a left-sided test tooth initially elicited an increase in band power in the contralateral (right) central parietal region (CP6). This activation then intensified and spread bilaterally, exhibiting an increase in band power across both the contralateral and ipsilateral somatosensory cortices in the final 1 second preceding the response. This aligns with previous studies that utilised functional near-infrared spectroscopy (fNIRS) to monitor brain activity while participants received cold stimulation on a sensitive tooth, both of which reported haemodynamic activation in the contralateral somatosensory cortex before participants signalled pain [22,23]

However, this does not fully explain our findings for the right-sided test tooth, which deviated by eliciting a predominantly ipsilateral (right-sided) increase in band power. All participants were instructed to use their left hand to press the ‘s’ key, which is an action neurologically driven predominantly by the right motor cortex. Since the motor execution of button presses by the left hand maps to the right hemisphere, this motor-related cortical activity spatially overlaps with the right central parietal region, an area known to be involved in sensorimotor integration and the planning of movement [24]. Additionally, because it is possible that some motor activity was not completely isolated from the cortical activity associated with nociception due to the aforementioned variations in motor execution, the increase in band power observed at the right central parietal region in both left- and right-sided test teeth likely reflects a combination of nociceptive processing and the motor preparation and execution of the button press [24]. This spatial overlap makes it challenging to definitively disentangle the nociceptive processing of discomfort from the motor response. To avoid this overlap in future studies, participants should be instructed to use the hand ipsilateral to the tooth being tested when pressing the button to indicate discomfort.

#### Comparison of EEG band power between response and no response trials

In addition to examining the spatial distribution of cortical activity in the seconds prior to participants indicating discomfort, comparing the magnitude of specific EEG frequency bands between response and no response trials provides insights into the neural mechanisms underlying the discomfort felt by participants. During the 2-second period preceding trial termination, there was a frequency-specific enhancement in delta band power (a low-frequency brain oscillation that occurs within the 1-4 Hz range) in the response trials across all four analysed channels: contralateral and ipsilateral somatosensory cortices, Cz, and Fz. These findings align with previous studies that have similarly reported significant increases in delta power in the somatosensory cortex [25] and frontal region (Fz) [26] after participants immersed their hands into iced water. The increase in these slow oscillations during cold stimulation is thought to reflect a mechanism of cortical inhibition directed at incoming information from afferent neurons [27]. Therefore, it may be that the enhanced delta power activity observed across the regions represents the brain’s attempt to dampen the perception of noxious nociceptive input [25] during cold stimulation of the tooth.

#### Correlation between peak-to-peak amplitude and measures of dentine hypersensitivity

To investigate the suitability of peak-to-peak amplitude as an objective marker for dentine hypersensitivity, correlation analysis was performed between the amplitudes and measures of dentine hypersensitivity. A significant correlation was found between the peak-to-peak amplitude at Fz and both the VAS score and DHEQ score. This is consistent with findings from a previous study, which also found a significant correlation between the peak-to-peak amplitude at Fz and the participants’ perceived intensity of thermal laser stimulation to the back of their hand [28]. The strong correlation between peak-to-peak amplitude at Fz with the VAS score and DHEQ score may therefore indicate that it can serve as an indicator of the participants’ perceived intensity of sensation. However, since Fz overlies the central frontal region and captures activity related to motor planning, as well as attention and cognitive control [10], peak-to-peak amplitude at Fz may also be a potential marker for attentiveness. Supporting this, Beydoun et al. [28] also linked the peak-to peak amplitude at Fz with levels of attention, showing that the amplitude was reduced when the participant’s attention was diverted.

Furthermore, the peak-to-peak amplitude exhibited a significant correlation with the temperature at which participants indicated discomfort across the ipsilateral and contralateral somatosensory cortices, as well as at Fz. The temperature of response effectively represents the participant’s threshold for the cold stimulus. These correlations suggest that participants who reached their threshold of discomfort at higher temperatures had a higher degree of synchronous neural firing [29] across networks responsible for processing and evaluating sensory information.

## Conclusion

To the best of our knowledge, this is the first study to record EEG activity during tooth stimulation. While further experiments are needed to refine the experimental paradigms in order to improve the quality of EEG data collected, this study provides strong evidence that components of EEG responses to evaporative and thermal stimulation could serve as objective measures of the severity of dentine hypersensitivity.

## Acknowledgments

We would like to thank all participants for taking part in this study. We gratefully acknowledge the staff at the Charles Clifford Dental Hospital and the School of Clinical Dentistry, University of Sheffield, UK, who helped with the recruitment of participants and administration of questionnaires, in particular Katy D’Apice.

## Author contributions

FMB, MA, MWB and CRP designed and supervised the study. NW and HB carried out the study. NW analysed the data and drafted the manuscript. FMB, MA, MWB and CRP reviewed and edited the first draft. All authors reviewed and approved the final paper.

